# Transcription and replication organize cohesin-dependent chromosome loops

**DOI:** 10.1101/2020.12.16.423068

**Authors:** Kristian Jeppsson, Toyonori Sakata, Ryuichiro Nakato, Stefina Milanova, Katsuhiko Shirahige, Camilla Björkegren

## Abstract

Genome function and stability is strictly dependent on regulated folding of chromosomes in space and time. Chromosome loop formation by the protein complex cohesin is a central feature of this multilayer organization^1^. Accumulating evidence indicates that cohesin creates loops by extruding chromosomal DNA through its ring-like structure in a process that is controlled by the cohesin loading factor Scc2 and the unloader Wpl1^1–4^. Cohesin’s chromosomal positioning is affected by transcription in both yeast and human cells^5,6^, and the complex localizes in the vicinity of replication forks^7^. However, if transcription directly influences chromosome looping remains an unresolved question, and the threedimensional organization of replicating chromosomes is unknown. Here we show that transcription and replication machineries create chromosome loop boundaries. We find that drug-induced depletion of chromosome-bound RNA polymerases triggers a rapid expansion of cohesin-dependent chromosome loops in the budding yeast *Saccharomyces cerevisiae.* New loop boundaries also form at a few highly expressed genes induced by the cellular stress response caused by the drug. The results also reveal that S-phase chromosomes are shaped by cohesin-dependent loops organized by transcription, with additional loop anchors at replication forks. Together, our results show that replication and transcription control the three-dimensional organization of the genome by blocking the progression of loop-forming cohesin. They also open for the possibility that the resulting positioning of loop-forming cohesin in the vicinity of transcription and replication machineries is part of cohesin’s functions in transcription control, sister chromatid cohesion and maintenance of fork stability.

## Transcription inhibition removes boundaries for cohesin-dependent chromosome loops

Cohesin belongs to the family of structural maintenance of chromosome (SMC) protein complexes and was initially identified as a tether of sister chromatids, the products of chromosome replication^8,9^. This cohesion, established in close vicinity to the advancing replication fork and removed at anaphase onset, is essential for correct chromosome alignment and segregation during mitosis^10–12^. Cohesin complexes involved in sister chromatid cohesion are stably associated to chromosomes^13^, and insensitive to inhibition of the cohesin loading factor Scc2 (NIPBL)^14^. In addition to being localized at centromeres, these stable complexes are found in between convergently oriented genes in *S. cerevisiae^6^.* This positioning is generally thought to reflect how transcribing RNA polymerases push cohesin into place. Later investigations using genome-wide chromosome conformation capture (3C) techniques have revealed that cohesin also creates chromosome loops^1,3^. Work in mammalian and insect cells show that interphase chromosomes are organized into so called topologically associated domains (TADs), which are large chromosomal regions within which chromosome *cis* interactions occur more frequently than with neighbouring regions^15–17^. Current data indicates that *cis* interactions within a TAD can be largely explained by dynamic loop extrusion by cohesin^1,2^, and that the insulator protein CTCF plays an essential role in TAD boundary formation in mammalian cells^15,18^. More recent analyses show that *S. cerevisiae* sister chromatids are also shaped by cohesin into TAD-like structures, which are considerably smaller than mammalian TADs^19,20^. The boundaries of these TAD-like structures in *S. cerevisiae* are found at the previously identified cohesin binding sites between convergently transcribed genes and at centromeres^19–21^. Thus, the loops within the yeast TAD-like structures appear to be formed by a sub-fraction of loop-extruding, dynamic cohesin complexes which occasionally reach the boundaries of the TAD-like structure, and thereby link two proximal cohesin binding sites. Cohesin-dependent TAD-like structures with boundaries between convergently oriented genes also form on unreplicated *S. cerevisiae* chromosomes^19,20^, which shows that the boundaries for loop expansion are independent of sister chromatid cohesion. This opens for the possibility that the boundaries are created by, or as a consequence of, transcription.

To test if transcription controls chromosome loop boundaries, we used genome-wide Hi-C analysis to determine how thiolutin, an inhibitor of yeast RNA polymerases^22^, influences chromosome structure in G2/M-arrested *S. cerevisiae* cells. Initial ChIP-sequencing (ChIP-seq) and quantitative ChIP-PCR (ChIP-qPCR) analysis of the RNA polymerase II (RNA pol II) subunit Rpo21 confirmed a reduction of RNA pol II in most open reading frames 30 minutes after addition of the drug (Fig. S1a-c). Subsequent Hi-C experiments showed that TAD-like structures were substantially reduced after the addition of thiolutin, while new long-range *cis* interactions appeared (Fig. 1a-d, S2a-c, S8a). More specifically, *cis* interactions ranging between ≈ 3 to 30 kb decreased, while those ≈ 40 kb to > 500 kb strongly increased when transcription was inhibited (Fig. 1d). Owing to weak residual chromosome loop barriers at cohesin sites in the thiolutin-treated cells (Fig. 1a, S8a), it was also possible to perform aggregate peak analysis. This reveals the aggregated contact signal between pairs of cohesin binding sites separated by increasing distances and normalized to random sites^19^, and the results confirmed a thiolutin-induced increase in chromosome loop size (Fig. S4a). The TAD-like structures were also drastically reduced after depletion of Scc2 in G2/M-arrested cells (Fig. 1a and c, S2a and c, S3a and e, and S4a, S8b), thereby confirming that G2/M-phase chromosome loops depend on cohesin loading. However, in contrast to the thiolutin-treated cells, no new long-range *cis* interactions were observed in the contact maps after depletion of Scc2 (Fig. 1a-d, S2a-c, S4a). To compare the effect of thiolutin to the reported increase in average loop size in cells lacking the cohesin unloading factor Wpl1 (Rad61/WAPL)^19^, we performed Hi-C analysis after G2/M-specific depletion of Wpl1 (Fig. S3a and e). This revealed the expected increase in loop size upon Wpl1 depletion (Fig. 1a-d, S2a-c, S4a, S8b). The results also showed that in contrast to thiolutin-treated cells, the TAD-like structures detected in Wpl1-depleted cells are more clearly limited by cohesin binding sites (Fig. 1a, S4a, S8b). Contact probability plots also showed that the change in loop size is less dramatic than after thiolutin treatment (Fig. 1d). Wpl1-depletion also changes interactions in a more limited extent in terms of loop size, only causing an increase of interactions ranging between ≈ 40 kb to ≈300 kb (Fig. 1d). In further support for transcription inhibition removing chromosome loop barriers in contrast to depleting chromosome-bound cohesin, we found that centromeres still prevent interactions between arms on the same chromosomes in thiolutin-treated cells, while this barrier is weakened after Scc2 depletion (Fig. S2a and c). Moreover, ratio plots of all 16 *S. cerevisiae* chromosomes show that thiolutin increases chromosome *cis* interactions and decrease *trans* interactions, which is similar to changes caused by Wpl1-depletion, but opposite to what is detected after Scc2-depletion (Fig. 1c, right-hand panels).

**Figure 1.**
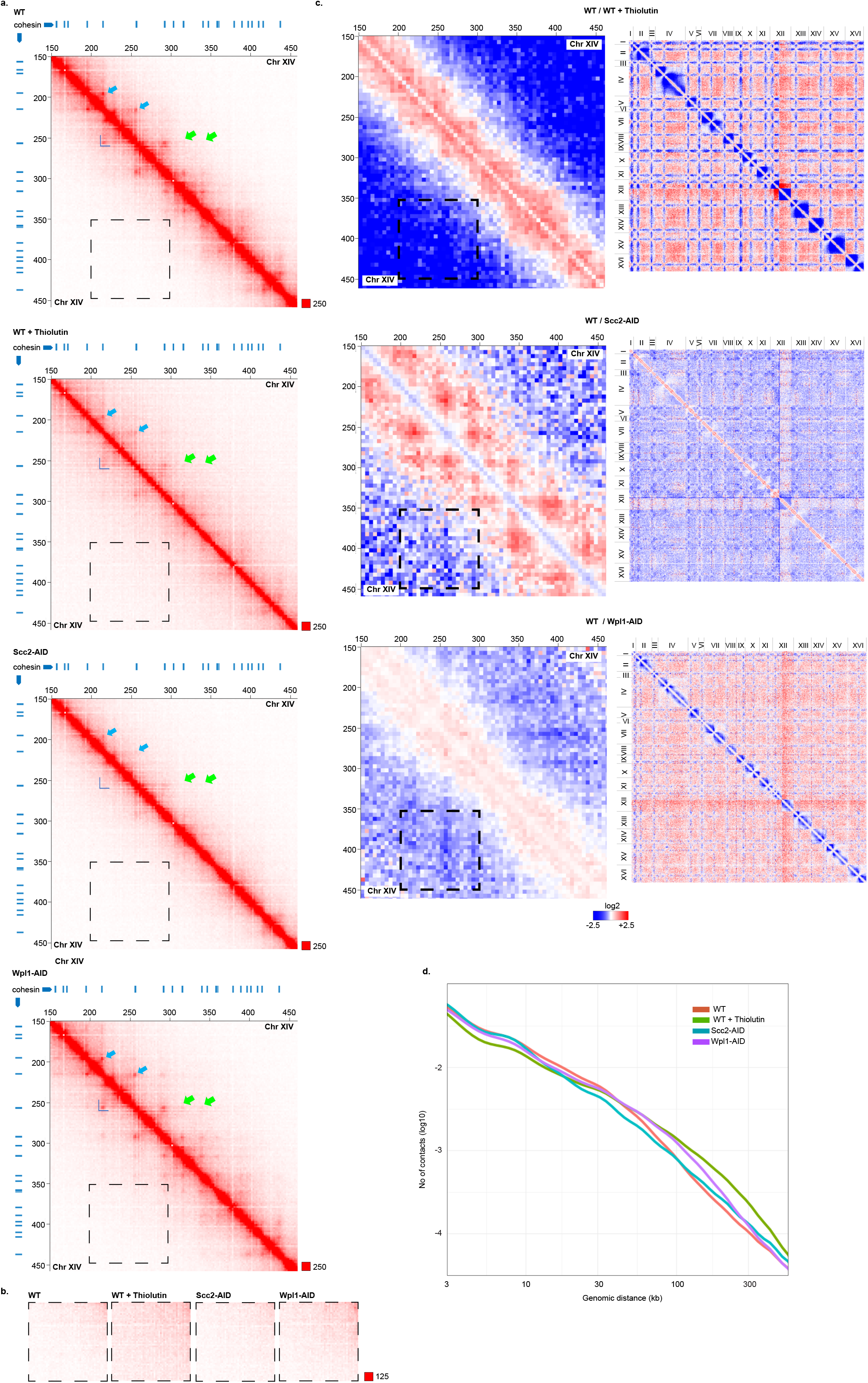
Transcription inhibition removes chromosome loop barriers, triggering the formation of long-range *cis* interactions. **a.** Normalized Hi-C contact maps (2 kb binning) showing *cis* interactions along the arm of chromosome XIV (150-450 kb from left telomere) in G2/M-arrested, untreated and thiolutin-treated wild type (WT) cells, or after Scc2 and Wpl1 depletion (Scc2-AID, Wpl1-AID) as indicated. Blue lines on top and to the left of the panels: cohesin binding sites, dark blue L-shape: TAD-like structure, light blue arrows: chromosome loop anchors, light green arrow: chromosome loop anchors detected after Wpl1 depletion. **b.** Normalized Hi-C contact maps (2 kb binning) in regions indicated by black, dashed squares in panels displayed in a), highlighting changes in long-range *cis* interactions. **c.** Normalized Hi-C ratio maps (without binning) comparing chromosome interactions in G2/M-arrested WT cells with thiolutin-treated, Scc2- or Wpl1-depleted cells as indicated. Left-hand panels: *cis* interaction ratios along the same chromosomal regions as depicted in a). Black, dashed squares indicate regions highlighted in a) and b). Right-hand panels: *cis* and *trans* interaction ratios along and between all 16 *S. cerevisiae* chromosomes. **d.** Contact probability plots as function of genomic distance (log10 scale) comparing interactions in G2/M-arrested untreated WT cells to thiolutin-treated, Scc2- or Wpl1-depleted cells as indicated. The experiments were performed using the growth protocol outlined in Figure S3a.

Together, this shows that transcription inhibition induces changes in chromosomes conformation that cannot be explained by lack of cohesin loading or unloading. Instead, the increase in loop size and percentage of *cis* interactions, and the maintenance of the centromere barrier, indicate that cohesin loading still occurs, but loop size is altered upon transcription inhibition. This indicates that transcription inhibition prevents loop boundary formation along chromosome arms and allows cohesin to extend “supersized” loops. If these loops are formed by cohesin, they should depend on Scc2 and be stabilized by Wpl1-depletion. Hi-C analysis of thiolutin-treated G2/M-arrested cells that had been depleted of Scc2 or Wpl1 before addition of the drug showed that this is indeed the case (Fig. S4b-c).

## Enrichment of dynamic cohesin between convergently expressed genes on unreplicated chromosomes depends on transcription

If transcription inhibition allows cohesin to extrude loops beyond normal boundaries, cohesin accumulation between convergent genes is expected to diminish. Thiolutin treatment has indeed been reported to reduce cohesin accumulation at some sites along chromosome arms in G2/M-arrested cells^20^. Revisiting this, we analysed the binding pattern of the cohesin subunit Scc1 (RAD21) by normalized ChIP-seq (see methods) and ChIP-qPCR, with or without prior addition of thiolutin. This provided similar results as in^20^, *i. e.* minor reduction of cohesin enrichment at some chromosome arm sites and no alterations in the centromeric regions after thiolutin treatment (Fig. S5a-c). Average peak plots of Scc1 binding sites along chromosome arms revealed a general broadening of peaks (Fig. S5b), reflecting a thiolutin-induced splitting of Scc1 peaks into two (Fig. S5a), and indicating a change in positioning in response to transcription inhibition. Even if these alterations are in line with a reorganization of dynamic cohesin in response to removal of chromosome loop barriers, the interpretation is uncertain due to the presence of stably associated cohesive cohesin complexes. To enable analysis of dynamic cohesin complexes alone, we investigated the effect of thiolutin on the localization of Scc1 in cells arrested in early S-phase by addition of hydroxyurea (HU), in which replicated and unreplicated regions have been mapped with high precision (see for example^23^). In the unreplicated regions, which are devoid of stable cohesive cohesin complexes, thiolutin treatment led to a general and significant reduction of Scc1 enrichment in intergenic regions between convergently expressed genes (Fig. 2a-c, S3b). In contrast, ChIP-qPCR analysis showed that cohesin levels increase at an unreplicated “non-convergent” site (Fig. 2c, *BPH1* ORF). These results validate the Hi-C analysis which indicates that transcription inhibition removes barriers that block the progression of dynamic loop-extruding cohesin (Fig. 1, S2 and S4), thereby reducing cohesin accumulation between convergent genes, and increasing the levels in other regions of the chromosomes.

**Figure 2.**
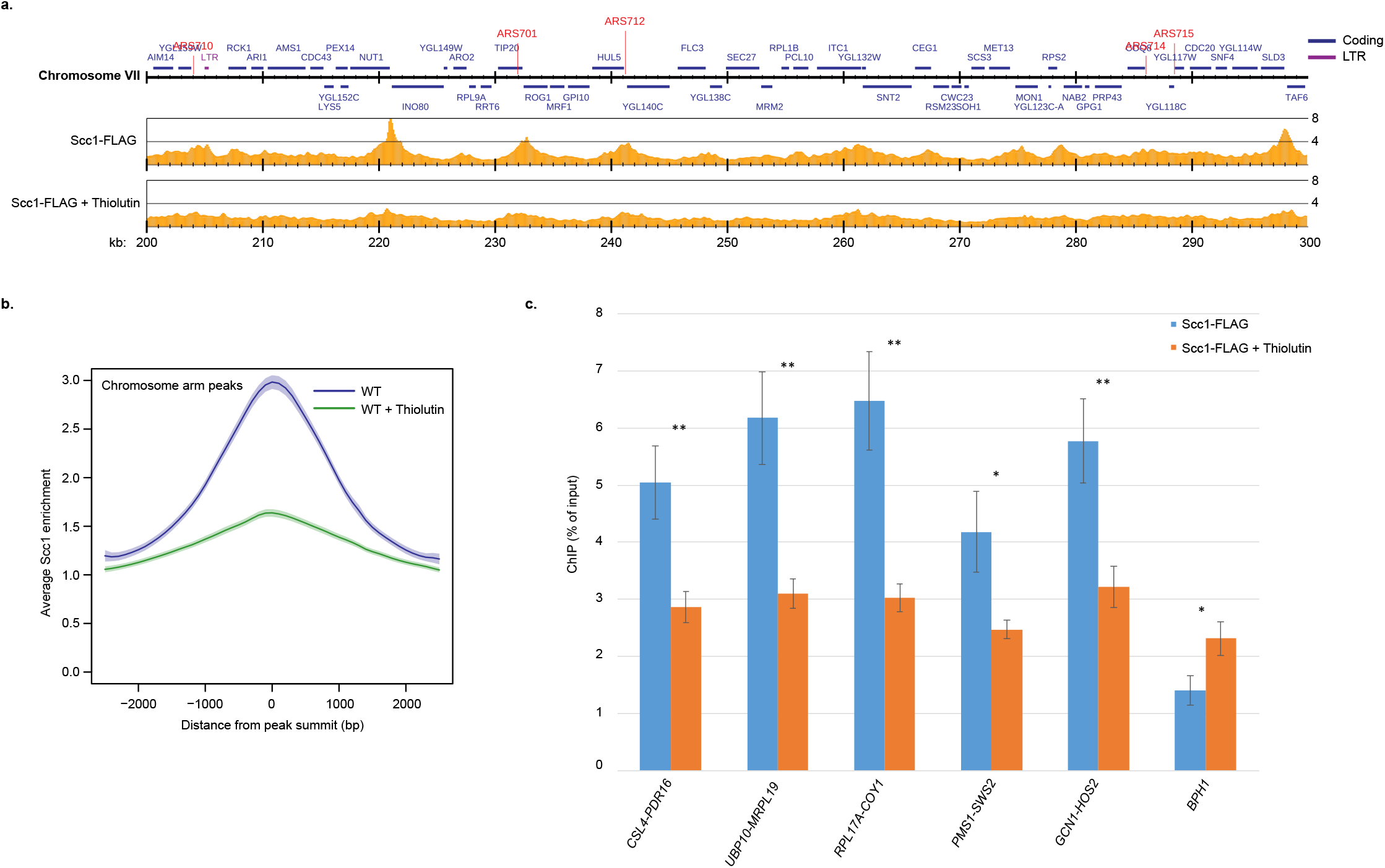
Transcription inhibition removes dynamic cohesin from its normal localization between convergently oriented gene pairs. **a.** Map of Scc1-FLAG enrichment in an unreplicated region of chromosome VII (200-300 kb from left telomere) based on normalized ChIP-seq analysis of untreated and thiolutin-treated wild type (WT) cells arrested in S-phase. The Y-axis shows fold enrichment of ChIP / input in linear scale, the X-axis shows chromosomal positions. Blue and purple horizontal bars in the uppermost genomic region panel denote coding regions and long terminal repeats (LTRs), respectively, red vertical lines indicate replication origins (ARS). **b.** Average peak plots of Scc1-FLAG binding sites along chromosome arms, based on the analysis presented in a). **c**. Chromosomal association of Scc1-FLAG at selected intergenic regions (indicated by flanking gene-pairs), and within the *BPH1* ORF, as determined by ChIP-qPCR of samples collected from cells treated as in a). *N=3, *: p≥0.05, **: p≥0.01.* The experiments were performed using the growth protocol outlined in Figure S3b.

## Highly expressed genes and replication forks hinder cohesin-dependent loop expansion

When inspecting the contact maps obtained from thiolutin-treated cells, a low number of new boundaries were detected along chromosome arms. These were generally found at stress response genes on which RNA pol II levels increased in response to thiolutin (Fig. 3a-b, S6a-c, S8c), which contrasts with most open reading frames (Fig. S1), but agrees with the reported cellular stress response induced by the drug^24^. The *cis* interactions anchored at stress response genes were weakened by depletion of Scc2 before thiolutin treatment, and strengthened, displaying stronger loop anchor signals, when Wpl1 was pre-depleted (Fig. 3b). Scc2- and Wpl1-regulated, thiolutin-induced boundary formation at these sites was also confirmed by insulation score analysis (Fig. 3c). Together, this provides additional evidence that the transcription machinery inhibits the progression of loop-forming cohesin, thereby controlling chromosome loop organization.

**Figure 3.**
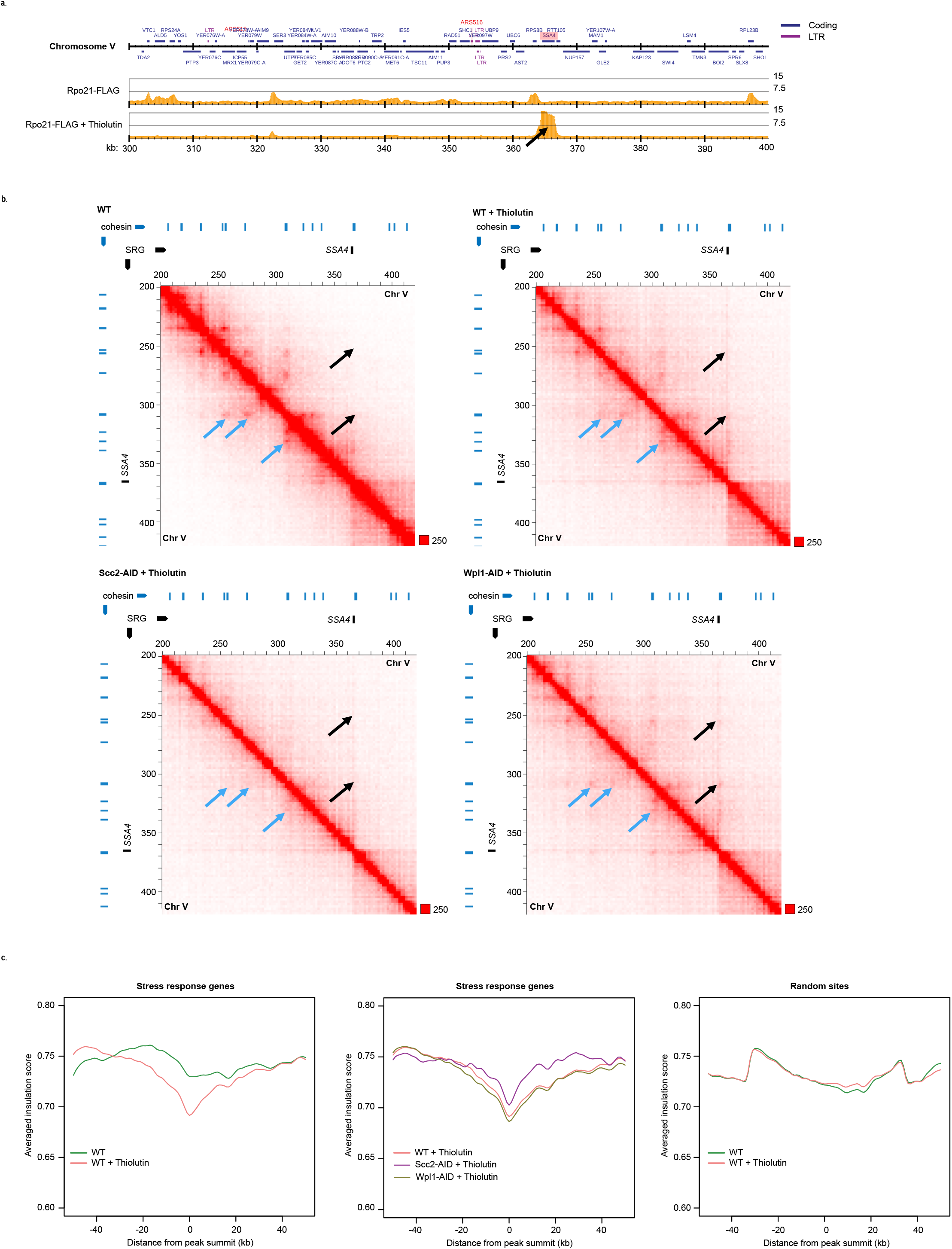
Highly expressed genes create novel chromosome loop barriers. **a.** Map showing Rpo21-FLAG enrichment on part of chromosome V, (300-400 kb from left telomere) in the absence or presence of thiolutin, based on normalized ChIP-seq analysis. Annotations as in Figure 2. Black arrow indicates thiolutin-induced Rpo21-FLAG accumulation at *SSA4.* The experiment was performed using the growth protocol outlined in Figure S3c. **b.** Normalized Hi-C contact maps (2 kb binning) showing *cis* interactions along the arm of chromosome V (2004-20 kb from left telomere) in G2/M-arrested untreated and thiolutin-treated wild type (WT), Scc2- and Wpl1-depleted cells (Scc2-AID, Wpl1-AID). Black arrows indicate barrier formation and *cis* interactions at the *SSA4* gene. Blue arrows highlight interactions that are reduced by thiolutin treatment. Blue and black lines on top and to the left of the panels: cohesin binding sites and stress response gene (SRG), respectively. **c**. Averaged insulation score plots based on normalized Hi-C data from untreated and thiolutin-treated G2/M-arrested wild type (WT) cells, and thiolutin-treated, Scc2-or Wpl1-depleted (Scc2-AID, Wpl1-AID) cells. The analysis focuses on 100 kb regions spanning 35 stress response genes that display thiolutin-induced increase in Rpo21-FLAG association (left-hand and middle panels), or 35 random sites (righthand panel). The experiments in b) and c) were performed using the growth protocol outlined in Figure S3a.

The observation that thiolutin reduces cohesin accumulation between convergent genes on replicating chromosomes suggest that cohesin-dependent loops also form during S-phase (Fig. Fig. 2a-c). Hi-C analysis of HU-arrested wild type, Scc2- and Wpl1-depleted cells revealed that this is indeed the case (Fig. 4a, S7a). In addition to revealing TAD-like structures flanked by cohesin-binding sites between convergently oriented gene pairs, the resulting contact maps disclosed strong loop barriers at replication forks (Fig. 4a, S7a). This indicates that loopextruding cohesin is blocked by the replication machinery. This was supported by aggregate peak analysis focusing on all early replication origins along chromosome arms (Fig. S7b, three upper panels). The analysis also revealed Scc2-dependent interactions between replication forks and cohesin sites that are stabilized in Wpl1-depleted cells (Fig. S7b, four lower panels). Together, this shows that replication and transcription machineries control cohesin-dependent chromosome looping during S-phase.

**Figure 4.**
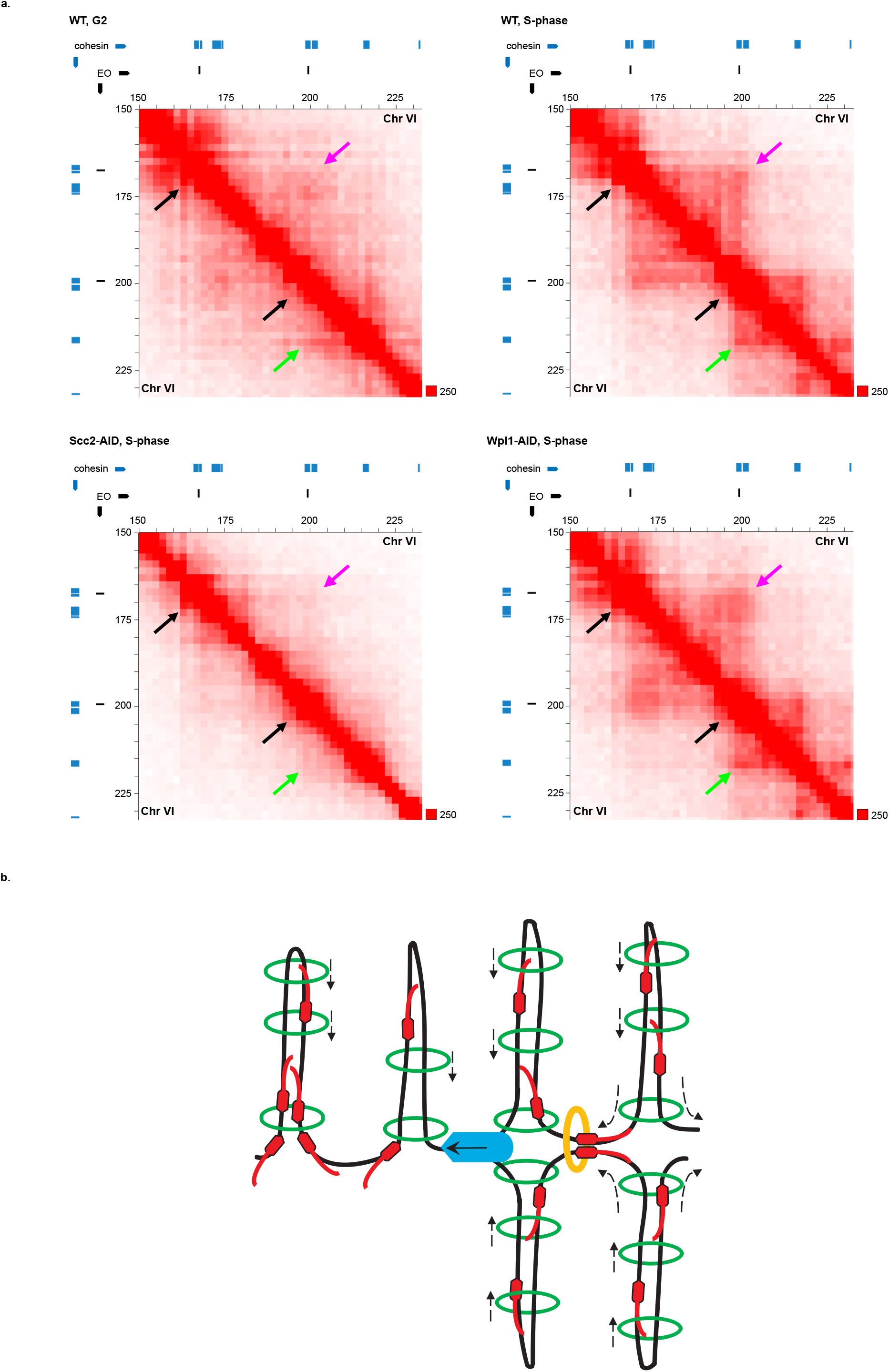
Replication forks act as chromosome loop barriers. **a.** Normalized Hi-C contact maps (2 kb binning) showing *cis* interactions along the arm of chromosome VI (150-230 kb from left telomere) in G2/M-arrested wild type (WT) cells, and S-phase-arrested WT, Scc2- or Wpl1-depleted (Scc2-AID, Wpl1-AID) cells. Black arrows highlight loop boundaries formed at origins and weakened after Scc2 depletion. Green arrows indicate a replication origin-cohesin site interaction that appears in S-phase and is weakened by Scc2 depletion. Pink arrows indicate S-phase specific loop anchor, weakened by Scc2 depletion and strengthened by Wpl 1 depletion. Blue and black lines on top and to the left of the panels: cohesin binding sites and early firing origin (EO), respectively. The experiments were performed using the growth protocol outlined in Figure S3c (G2/M) and S3d (S-phase). **b**. Schematic model of obtained results. Loop extruding cohesin complexes (green open circles) are blocked by head-on collisions with transcribing RNA polymerases (red, pointed ovals with “RNA-tail”), which thereby create loop barriers on unreplicated and replicated chromosomes. The replication machinery (clear blue, pointed oval) also creates a barrier by blocking loop-extruding cohesin complexes. The resulting positioning of cohesin complexes in the vicinity of polymerases might underlie cohesin’s functions in transcription regulation, fork stability and formation of cohesive complexes (orange open circle) in the wake of the fork. Dashed arrows indicate the direction of cohesin movement along chromosomes during extrusion.

Taken together, the presented results reveal an unexpected interplay between transcription, replication, chromosome three-dimensional organization and cohesin functions (Fig. 4b). Cohesin’s role in chromosome looping^1–3^, and transcription-dependent localization of the complex in both yeast and human cells are well established^5,6,25^. However, the relationship between these two features has previously not been directly addressed, and the current model, originating from analyses of stably bound, cohesive complexes, proposes that transcription pushes cohesin along chromosomes^6^. Even if this might still be true for complexes involved in sister chromatid cohesion, our analysis indicates that transcription-dependent positioning of dynamic cohesin complexes reflects a barrier function for transcription. This implies that the positioning of cohesin at 3’ends of convergently expressed genes is caused by head-on collisions between the convergently oriented transcription units and cohesin complexes moving into the intergenic region from opposite sides (Fig. 4b). The results also open for that stably bound cohesive complexes in between convergently oriented genes are pushed in place by loopforming cohesin and not the transcription machinery itself. The observation that bacterial SMC complexes which translocate along newly replicated chromosomes are blocked by convergently oriented genes^26,27^, indicates that the here observed cohesin-transcription interplay reflects an evolutionary conserved feature of SMC complexes. The evolutionary conservation of transcription as a roadblock for loop-forming cohesin is further supported by investigations showing that TAD boundaries correlate with active transcription in many species (reviewed in^28^). This said, the overall effect of transcription on chromosome loop formation is expected to vary depending on the presence of other barriers, such as the CTCF insulator protein^15,16,29^. The recently published results showing that cohesin is required for increased tethering of CTCF sites in response to transcription inhibition strengthen the notion that transcription can act in parallel with CTCF-dependent loop control^30^.

In addition to revealing a function for transcription in loop formation, our results also suggest that the cohesin function in gene regulation reflects a direct association with transcription units, and not only the function of the complex in chromosome looping. The observation that replication forks are barriers for loop-extruding cohesin complexes also have several implications. First, it indicates that previously reported accumulation of cohesin at replication forks reflects how loop-extruding complexes slide into place instead of being recruited from a soluble fraction (Fig. 4b)^7^. This opens for new models for S-phase specific functions of cohesin, including establishment of sister chromatid cohesion, which could depend on loop-extruding cohesins that are converted to cohesive complexes behind the fork (Fig. 4b). Second, cohesin might not only work on newly formed chromatids, but also in front of the fork, where the loopforming activity could influence fork progression and stability^7,31^. Third, a role for the replication machinery in determining chromosome looping during S-phase could be an important aspect in the establishment of TADs, and provides a potential mechanism for transcription-independent TAD formation during early development^32^. Altogether, the revelation that both transcription and replication machineries are barriers for cohesin-dependent loop expansion sets the stage for future analysis of chromosome organization and cohesin function from new perspectives.

## Acknowledgements

This study was supported by funding from Swedish Cancer foundation, Swedish research council and Centre for Innovative Medicine (CIMED) to CB, and JST CREST (JPMJCR18S5), Grant-in-Aid for Scientific Research(S) from JSPS (20H05686), and Grant-in-Aid for Transformative Research Areas from MEXT (20H05940) to KS. KJ was funded by JSPS Postdoctoral Fellowship for Overseas Researchers, and travel grants from The Scandinavia-Japan Sasakawa Foundation, Helge Ax:son Johnsons Stiftelse and a KI-UTokyo collaboration.

## Author contributions

KJ made the initial discoveries, KJ and CB developed the concept, and CB wrote and assembled the manuscript with input from all co-authors. KJ performed Hi-C and ChIP-seq in a collaborative effort between CB and KS laboratories. KJ and SM performed ChIP-qPCR and Western blots, and SM performed the FACS analysis. TS, RN and KS performed and developed the bioinformatic analyses in collaboration with KJ.

## Materials & Correspondence

Correspondence and request for materials can be directed to Kristian Jeppsson and Camilla Björkegren.

## METHODS

### Yeast strains, growth conditions, protein degradation and transcription inhibition

All strains are of W303 origin with the modifications listed in Table S1. Cells were cultured in YEP medium (1 % yeast extract, 2 % peptone, 40 μg ml^-1^ adenine) supplemented with 2 % glucose (YEPD). For arrest in G2/M, benomyl (Sigma, 381586)-containing YEPD medium was added to cells growing logarithmically for a final concentration of 80 μg ml^-1^. Cell cultures were then incubated for 90 minutes at 30°C achieving complete G2/M-arrest. For synchronization in S-phase, 3 μg ml^-1^ α-factor mating pheromone (Sigma, custom peptide WHWLQLKPGQPMY) was added every hour (a total of three additions) to cells growing logarithmically at 25°C. Upon complete G1-arrest, cells were released into medium containing 0.2 M hydroxyurea (HU) (Sigma, H8627), and S-phase was allowed to progress at 25°C. For transcription shut-off and degradation of Scc2 and Wpl1 in G2/M-arrest, auxin (3-indoleacetic acid, Sigma, I2886) and doxycycline (Sigma, D9891) were added for 1 hour at the final concentration of 1 mM and 5 μg ml^-1^, respectively. For transcription shut-off and degradation of Scc2 and Wpl1 in a synchronized S-phase, auxin and doxycycline were first added to G1-arrested cells for 30 minutes, and then for an additional hour in the HU-containing release medium at the same final concentrations as above. For transcription inhibition, thiolutin (Abcam, ab143556) was added to cell cultures for the final concentration of 20 μg ml^-1^ for 30 minutes. Cell cycle progression and arrests were confirmed using standard protocol for FACS analysis of ethanol-fixed, propidium iodide-stained cells (Fig. S3).

### Hi-C library preparation

Our Hi-C protocol was adapted for *S. cerevisiae* from^1^. Fifty ml of cell culture (at an OD of 1.0) was fixed with 3 % formaldehyde for 20 minutes at 30°C, before the reaction was quenched by adding glycine to 0.125 M final concentration for 5 minutes at room temperature. Cells were washed once with 1 x PBS, before being resuspended in 5 ml pre-spheroplasting buffer (100 mM PIPES (pH 9.4), 10 mM DTT). The cells were incubated 5 minutes at room temperature and then pelleted (1500 g for 5 minutes at room temperature), before being resuspended in 5 ml spheroplasting buffer (50 mM KH2PO4/K2HPO4 (pH 7.5), 1 M Sorbitol, 10 mM DTT). Twenty-five μl of 10 mg ml^-1^ 100T Zymolyase (Nacalai Tesque, 07665-55) was added and cells were incubated for 15 minutes at 30°C. The spheroplasts were pelleted (500 g for 5 minutes at 4°C) and washed twice with ice-cold spheroplasting buffer (containing only 1 mM DTT). The spheroplasts were then resuspended in 250 μl of ice-cold Hi-C lysis buffer (10 mM Tris-HCl (pH 8.0), 10 mM NaCl, 0.2 % Igepal CA630) supplemented with 50 μl 6 x Complete (Roche, 04693132001) and 3 μl protease inhibitor (Sigma, P8215), and incubated on ice for 15 minutes. The spheroplasts were pelleted (1500 g for 5 minutes at 4°C) and washed twice with 500 μl of ice-cold Hi-C lysis buffer. After the last wash step, the supernatant was discarded and the spheroplasts were incubated for 6 minutes at 62°C. Thereafter sodium dodecyl sulphate (SDS) was added to a final concentration of 0.2 % and the reaction was immediately and thoroughly mixed by inversion, and thereafter incubated at 62°C for 10 minutes. After addition of 80 μl H2O, 25 μl 10 % Triton X-100 was added to quench the SDS. The reaction was mixed by inversion and incubated at 37°C for 15 minutes. Thereafter, 28 μl 10 x NEB DpnII buffer and 500 units of DpnII (NEB, R0543M) was added, and the chromatin was digested overnight at 37°C. At the end of the incubation, the reaction was supplemented with 250 units of DpnII and incubated for 1 hour at 37°C. Then, the restriction enzyme was inactivated at 62°C for 20 minutes. The presence of intact and individual DNA masses throughout the spheroplasting, digestion and ligation steps was confirmed by DAPI (4’,6-diamidino-2-phenylindole)-staining and microscopy. Marking and repairing DNA ends, proximity ligation, crosslink reversal, DNA shearing, size selection, biotin pull-down, preparation for Illumina sequencing, final amplification (15 cycles) and purification was performed as in^1^. The Hi-C libraries were sequenced on Illumina HiSeq series with 150-bp paired-end sequencing according to the manufacturer’s recommendations.

### Hi-C data analysis

The Hi-C data were processed using Juicer with the default parameter set^2^. The sequenced reads were mapped to the *S. cerevisiae* genome obtained from Saccharomyces Genome Database (SGD) (http://www.yeastgenome.org/). The uniquely mapped read pairs were randomly resampled and arranged in the number of the lowest sample (55.9 million read pairs). See Table S3 for details. Contact matrices used for further analysis were coverage (sqrt)-normalized at 1kb and 2 kb resolution with Juicer. The matrices were visualized by Juicebox^3^. Intra-chromosomal contact frequency distribution was calculated using non-duplicated valid Hi-C contact pairs at genomic distances increasing by 1 kb.

### Aggregate peak analysis

Aggregated intensities of pixels corresponding to pairs of specific sites in the contact matrices were calculated using aggregate peak analysis (APA) with “-r 1000 -n 0 -w 20 -k VC_SQRT” option using juicertools version 1.9.9^1,2^. Briefly, APA calculates the sum of submatrices around paired genomic loci derived from the contact matrix. Each of these submatrices is a pixel square centred at a single pair of loci in the upper triangle of the contact matrix.

### Insulation score analysis

Insulation score analysis was performed as previously reported^4^. Briefly, the score was calculated for every 1 kb bin as the total number of contacts formed across the bin by pairs of loci located on the either side, up to 40 kb away using coverage (sqrt)-normalized contact matrices. The score was normalized by the mean of all insulation scores.

### Chromatin immunoprecipitation, qPCR and ChlP-seq library preparation

Chromatin immunoprecipitation was performed as previously described, with the following details and modifications^5^. One hundred ml of cell culture (at an OD of 1.0) was crosslinked with 1 % formaldehyde for 30 minutes at room temperature, followed by incubation at 4°C overnight. Chromatin was sheared to a size of 300-500 base pairs by sonication (Bandelin Sonopuls HD 2070.2) and IP reactions, with anti-FLAG antibody (Sigma, F1804) conjugated to Dynabeads Protein A (Invitrogen, 10002D), were allowed to proceed overnight at 4°C. After completing the immunoprecipitation and reversing crosslinks as in^5^, the DNA clean-up step was modified as follows. One hundred μl TE containing 10 μg RNase A (VWR, A3832) was added to IP and input fractions and the reactions were incubated for 1 hour at 37°C. Then, 2 μl of 50 mg ml^-1^ Proteinase K (Sigma, 000000003115879001) was added and the reactions were incubated for 2 hours at 37°C. Lastly, the DNA was purified using Qiagen PCR Purification kit according to standard instructions. For ChIP-qPCR, the quantitative PCR was performed using Fast SYBR Green (Applied Biosystems, 4385612) and primers listed in Table S2 on Applied Biosystem 7500 Real-Time PCR System according to the manufacturer’s instructions. The data were analyzed using two-sided *t*-test, and the presented graphs show mean values from biological triplicates with error bars representing standard deviation. For ChIP-seq, DNA from ChIP and input fractions was further sheared to an average size of approximately 150 bp by sonication (Covaris M220). Samples were then prepared for sequencing using the NEBNext Ultra II DNA Library Prep Kit for Illumina (NEB, E7645) according to the manufacture’s protocol. The libraries were sequenced using the HiSeq 2500 platform to generate single-end 65 bp reads. Sequenced reads were mapped to the *S. cerevisiae* genome using Bowtie2 version 2.4.1 with the default parameter set^6^. For BrdU-IP samples, we used previously published data^7^. The reads were downloaded from Sequence Read Archive (SRA) under accession IDs SRR1555037 and SRR1555038 and mapped them using Bowtie version 1.1.2 with “-n2 -k1” option because Bowtie2 does not allow colour-space fastq data. See Table S4 for details.

### ChIP-seq data analysis

To call peaks for Scc1, we identified bins in which the fold enrichment (ChIP / input) was more than 2.0. Peaks overlapping with long terminal repeats (LTRs) were excluded. To define cohesin sites on chromosome arms, Scc1 peaks that overlap with pericentromeric regions (25 kb spanning each centromere) were excluded. Stress response genes were defined as open reading frames with an average Rpo21 / input fold enrichment higher than 4.0 in the presence of thiolutin and lower than 2.0 in the DMSO control condition. To define replication origins that had fired under in HU-arrest *(i.e.* early origins), we obtained a list of all origins (autonomously replicating sequences, ARSs) in the *S. cerevisiae* genome from SGD (http://www.yeastgenome.org/), and selected ARSs overlapping with BrdU-IP peaks with a fold enrichment of more than 1.5. For early origins along chromosome arms, ARSs overlapping with pericentromeric regions (25 kb spanning each centromere) were excluded. To quantitatively compare the ChIP-seq peaks among samples, we normalized the ChIP-seq data using ChIP-qPCR. We calculated relative ratios of peak intensity among samples based on the 4~7 qPCR sites from common peaks (Table S5) and applied them as scaling factors for spike-in ChIP-seq normalization. We used DROMPAplus version 1.8. for normalizing, peak-calling and visualizing ChIP-seq data^8^.

### Protein extraction and western blot

To monitor the degradation of HA-tagged Wpl1-AID or Scc2-AID, protein extraction was performed using standard trichloroacetic acid (TCA)-precipitation or as in^9^, respectively. Western blot membranes were detected using anti-HA antibody, clone 12CA5 (Roche, 1666606).

## Data availability

The raw sequencing data and processed files for Hi-C and ChIP-seq will be available at Gene Expression Omnibus (GEO) under the accession number GSExx. All other data supporting the findings of this study are available in the manuscript and from the corresponding authors upon reasonable request.

---

**Figure S1, related to Figure 1.**
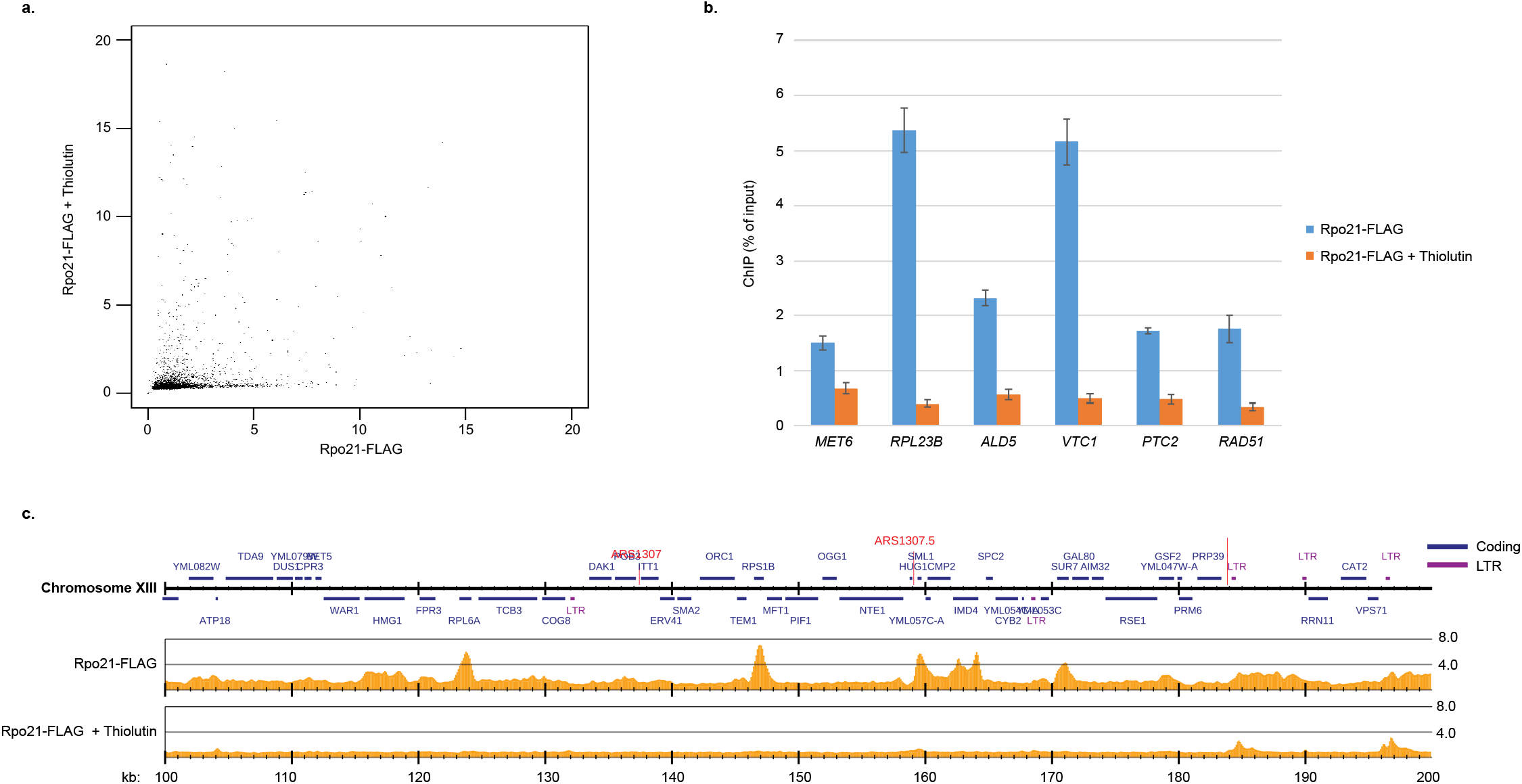
Thiolutin treatment reduces RNA pol II association in most open reading frames. **a.** Scatter plot comparing the accumulation of RNA pol II in 6570 *S. cerevisiae* open reading frames (ORFs) in the absence and presence of thiolutin based on normalized ChIP-seq analysis of Rpo21-FLAG in G2/M-arrested cells. Rpo21-levels were reduced in 93.5% of ORFs, increased in 6.5 %. **b.** Chromosome association of Rpo21-FLAG in indicated ORFs as revealed by ChIP-qPCR. N=3, p≤0.001 for all sites. **c.** Map of Rpo21-FLAG enrichment on chromosome XIII (100-200 kb from the left telomere) determined by normalized ChIP-seq. The Y-axis shows fold enrichment of ChIP / input in linear scale, the X-axis shows chromosomal positions. Blue and purple horizontal bars in the genomic region panel denote coding regions and long terminal repeats (LTRs), respectively, red horizontal lines indicate replication origins (ARS). The experiments were performed using the growth protocol outlined in Figure S3c.

**Figure S2, related to Figure 1.**
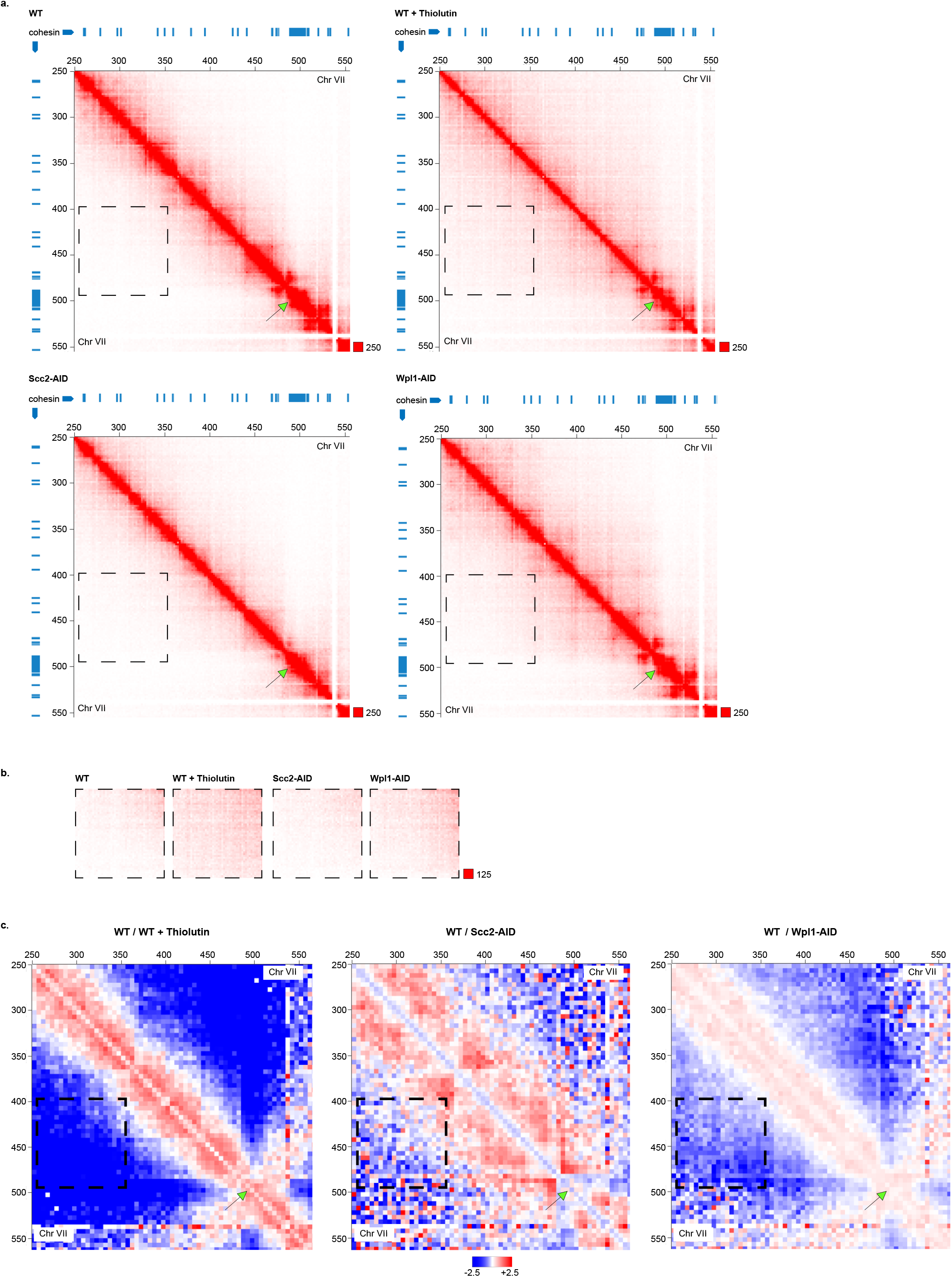
Transcription inhibition removes most chromosome loop barriers but leaves the centromere barrier function unperturbed. **a.** Normalized Hi-C contact maps (2 kb binning) showing *cis* interactions along a centromere-spanning region of chromosome VII (250-550 kb from left telomere) in G2/M-arrested, untreated and thiolutin-treated wild type (WT) cells, and after Scc2 and Wpl1 depletion (Scc2-AID, Wpl1-AID) as indicated. Blue lines on top and to the left of the panels: cohesin binding sites, black-rimmed green arrow: centromere VII. **b.** Normalized Hi-C contact maps (2 kb binning) in regions indicated by black, dashed squares in panels displayed in a), highlighting changes in long-range *cis* interactions. **c.** Normalized Hi-C ratio maps (no binning) comparing chromosome interactions in G2/M-arrested WT cells with thiolutin-treated WT, Scc2-or Wpl1-depleted cells as indicated, focusing on the same chromosomal region as a). Black-rimmed green arrow: centromere VII. The experiments were performed using the growth protocol outlined in Figure S3a.

**Figure S3, related to Figure 1-4.**
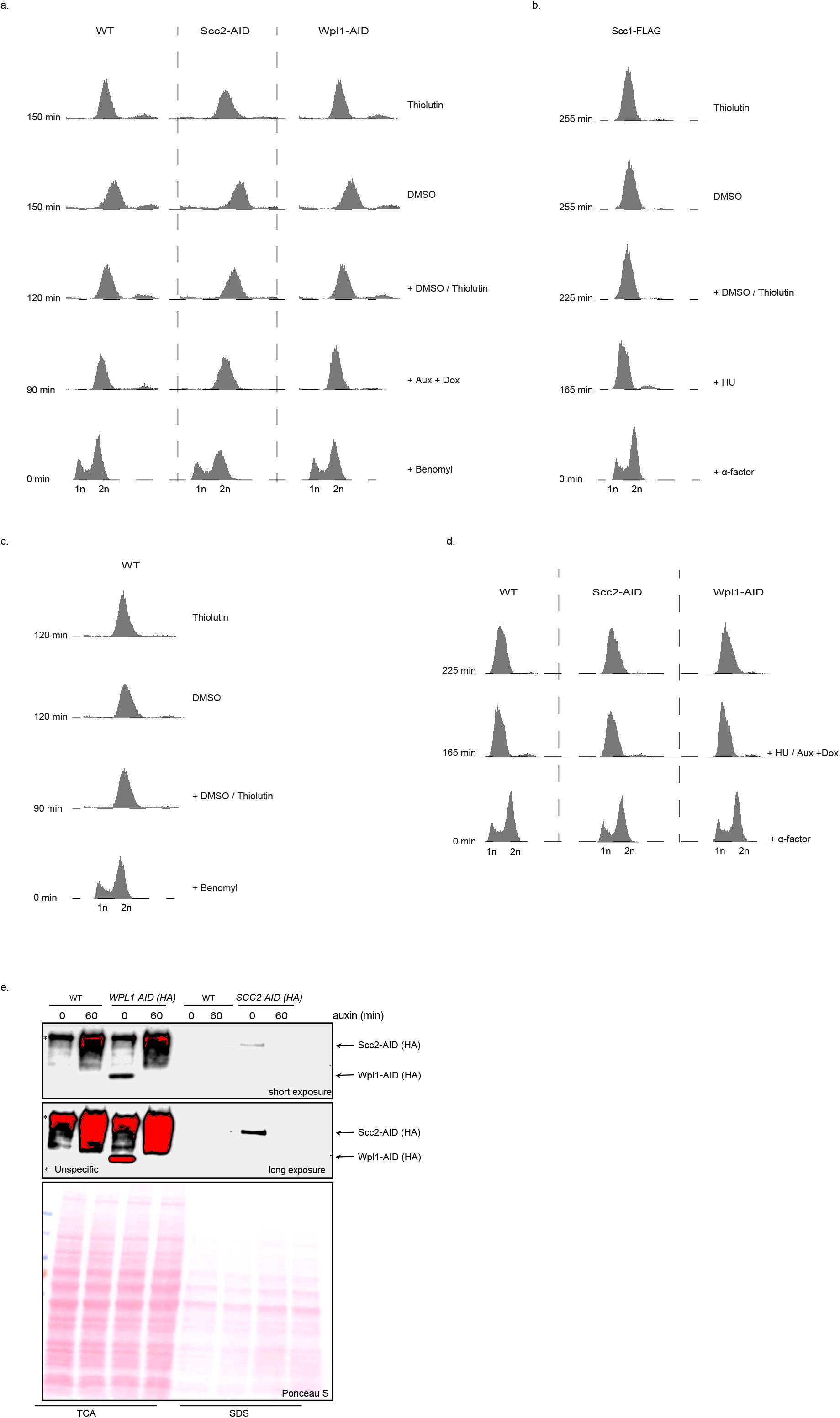
Representative analyses of cell cycle progression and Scc2 and Wpl1 depletion. **a.** Fluorescence-activated cell sorting (FACS) analysis of indicated cell types, samples for Hi-C, ChIP-seq and ChIP-qPCR were collected at the 150 minutes time point. Logarithmically growing cells were first arrested in G2/M by 90 minutes benomyl treatment, and subsequently treated with auxin and doxycycline for 30 minutes to deplete Scc2 or Wpl1. Thereafter, the cultures were split into two and treated with either DMSO or thiolutin for 30 minutes under maintained G2/M-arrest and Scc2- and Wpl1-depleting conditions. Experiments presented in Figures 1, 3b-c, S2, S4 and S6c were performed using this growth protocol. **b**. FACS analysis of wild type cells first arrested in G1 by addition of the alpha-factor pheromone for 165 minutes, and subsequently released into medium containing hydroxyurea to arrest cells in early S-phase. After 60 minutes, half of the culture was treated with either DMSO or thiolutin for 30 minutes when samples were collected. Experiments presented in Figure 2 were performed using this growth protocol. **c.** FACS analysis of logarithmically growing wild type cells initially arrested in G2/M by 90 minutes benomyl treatment. Thereafter the cultures were split in two and treated with either DMSO or thiolutin for 30 minutes under maintained G2/M-arrest, when samples for Hi-C and ChIP-seq analysis were collected. Experiments presented in Figures 3a, 4a (G2/M), S1, S5, S6a-b, S8a and S8c were performed using this growth protocol. **d.** FACS analysis of indicated cell types, first arrested in G1 by addition of the alpha-factor pheromone for 165 minutes. Thereafter, the cells were released into medium containing hydroxyurea to arrest cells in early S-phase (WT, Scc2- and Wpl1-AID), and auxin and doxycyline to deplete Scc2 or Wpl1 (Scc2-AID, Wpl1-AID). Samples for Hi-C analysis, ChIP-seq and -qPCR were collected after an additional 60 minutes. Results from S-phase arrested cells presented in Figures 4a and S7 (S-phase) were obtained using this growth protocol. **e**. Representative Western blots showing auxin- and doxycycline-induced degradation of Scc2- and Wpl1 – AID with a HA-epitope integrated at their N-terminus. Note that the samples for detection of Wpl1 and Scc2 were prepared using different protein extraction methods using trichloroacetic acid (TCA) or sodium dodecyl sulfate (SDS). Bottom panels show PonceauS-stained transfer-membranes demonstrating equal loading within each set of samples.

**Figure S4, related to Figure 1.**
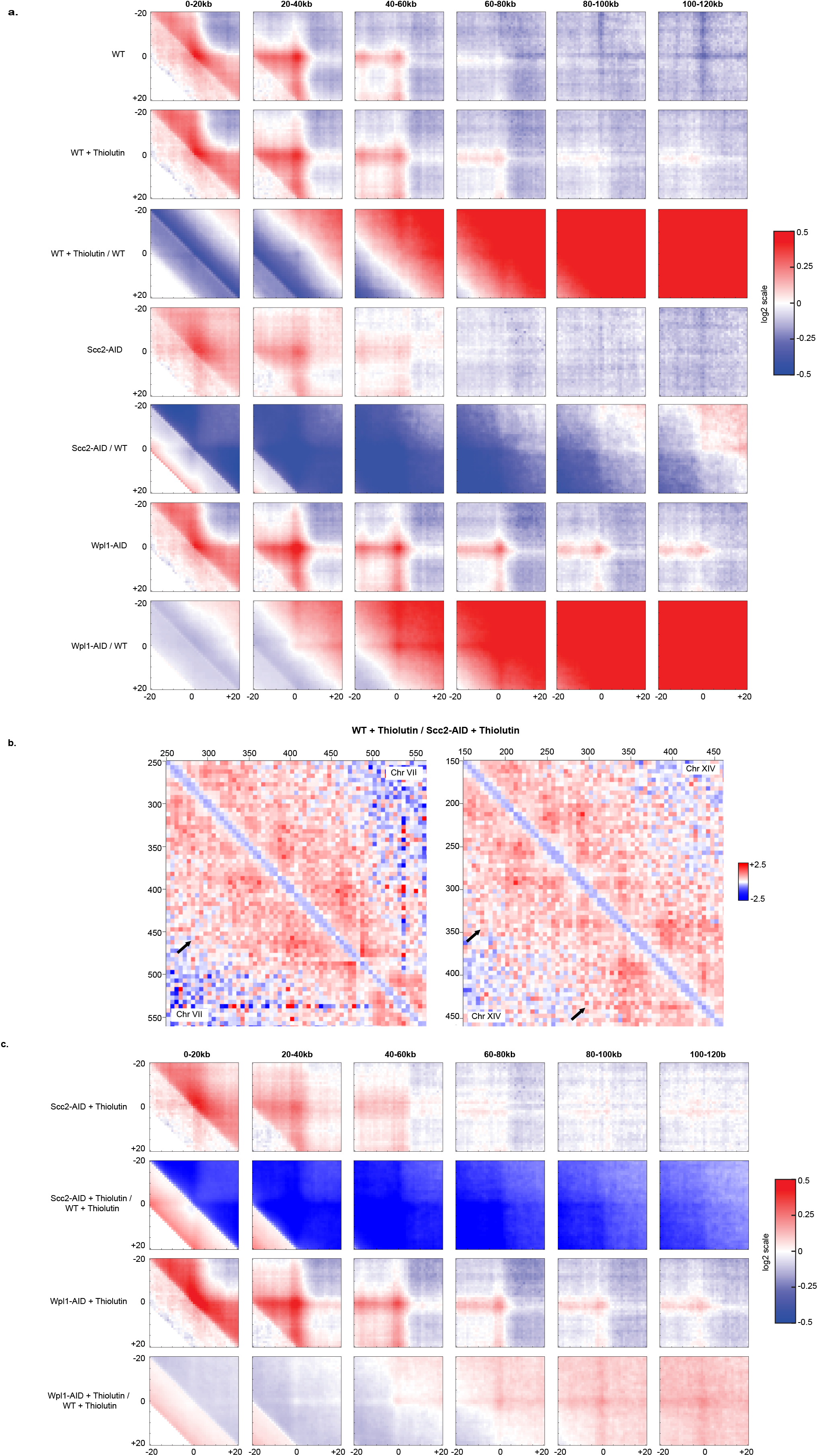
Transcription inhibition leads to novel long-range, cohesin-dependent *cis* interactions. **a.** Aggregate peak analysis centered on interactions between pairs of cohesin sites, separated by increasing chromosomal distances as indicated on top of the panels. The analysis is based on Hi-C analysis of G2/M-arrested, untreated and thiolutin-treated wild type (WT) cells, and cells depleted for Scc2 or Wpl1 (Scc2-AID, Wpl1-AID) during the G2/M-arrest. The interactions were either normalized to pairs of random sites (WT, WT+ Thiolutin, Scc2-AID or Wpl1-AID) or to cohesin-cohesin interactions obtained in WT cells (e.g. WT + Thiolutin / WT). Log2 color scale on the right-hand side. **b.** Normalized Hi-C ratio maps (without binning) comparing chromosome interactions in G2/M-arrested wild type (WT) and Scc2-depleted (Scc2-AID) cells, both treated with thiolutin. Black arrows indicate long-range chromosome interactions that predominate in WT, thiolutin-treated cells. Interactions ratios along the arm of chromosome VII (250-550 kb from left telomere, left-hand panel) and chromosome XIV (150-450 kb from left telomere, right hand panel) are shown. **c.** As in a) but performed on Scc2- and Wpl 1-depleted G2/M-arrested cells subsequently treated with thiolutin. Obtained interaction patterns are normalized to random sites or to interactions observed in G2/M-arrested WT cells treated with thiolutin. The experiments were performed using the growth protocol outlined in Figure S3a.

**Figure S5, related to Figure 2.**
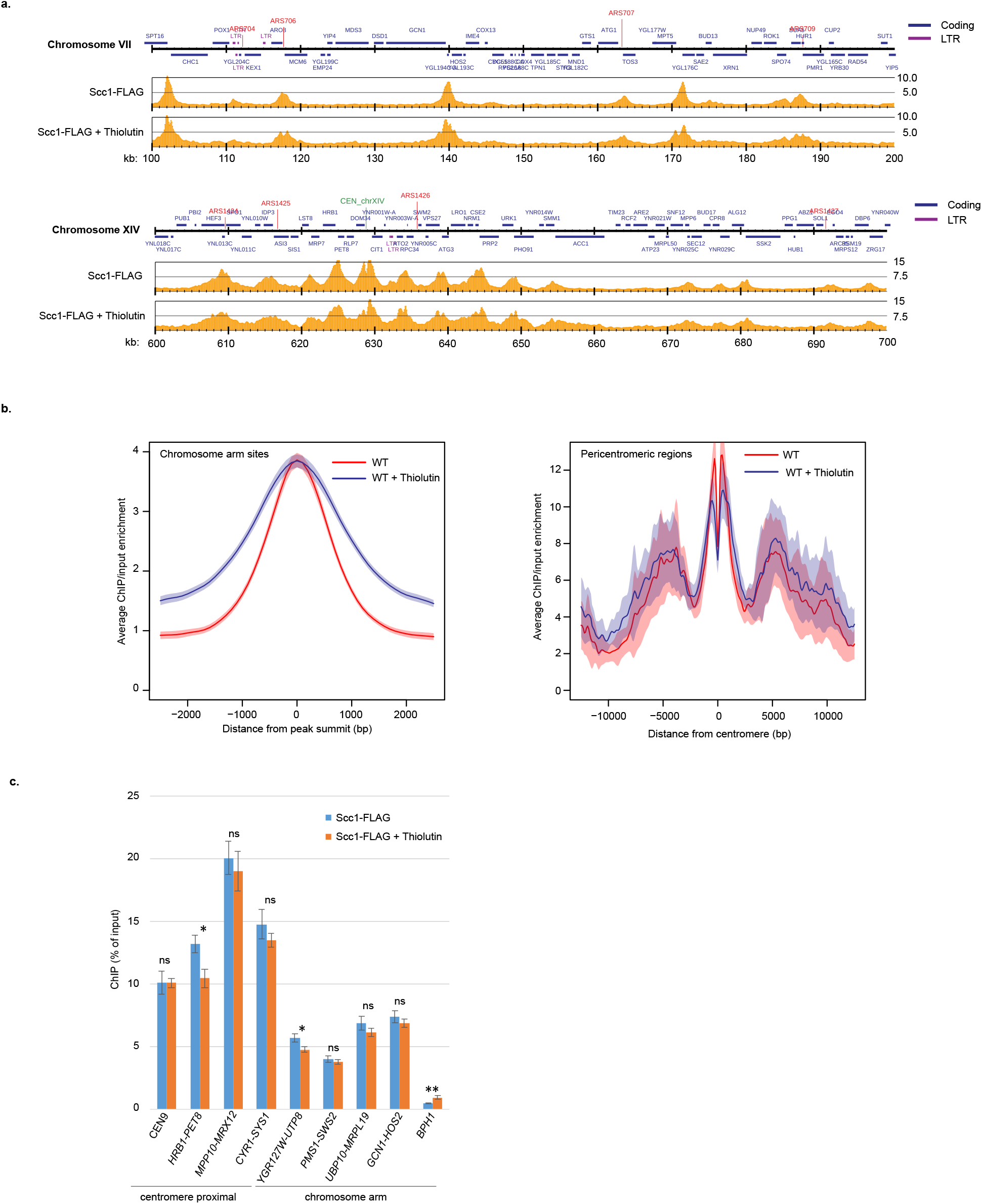
Transcription inhibition in G2/M leads to small changes in cohesin chromosomal localization. **a.** Maps based on normalized ChIP-seq analysis showing the enrichment of Scc1-FLAG on chromosome VII (100-200 kb from left telomere, upper map) and XIV (600-700 kb from left telomere, lower map) in untreated or thiolutin-treated wild type (WT) cells arrested in G2/M. The Y-axis shows fold enrichment of ChIP / input in linear scale, the X-axis shows chromosomal positions. Blue and purple horizontal bars in the uppermost genomic region panel denote coding regions and long terminal repeats (LTRs), respectively, red and green vertical lines indicate replication origins and centromeres, respectively (ARS, CEN). **b.** Average peak plots of all Scc1-FLAG binding sites along chromosome arms (left panel) and in 25 kb regions spanning each centromere, based on the analysis presented in a). **c**. Chromosome association of Scc1-FLAG at centromere IX, at selected intergenic regions flanked by indicated gene-pairs (centromere-proximal or along chromosome arms), and within the *BPH1* ORF, as determined by ChIP-qPCR of samples collected from cells treated as in a. *N=3, ns: p>0.05, *: p≥0.05, **: p≥0.01* The experiments were performed using the growth protocol outlined in Figure S3c.

**Figure S6, related to Figure 3.**
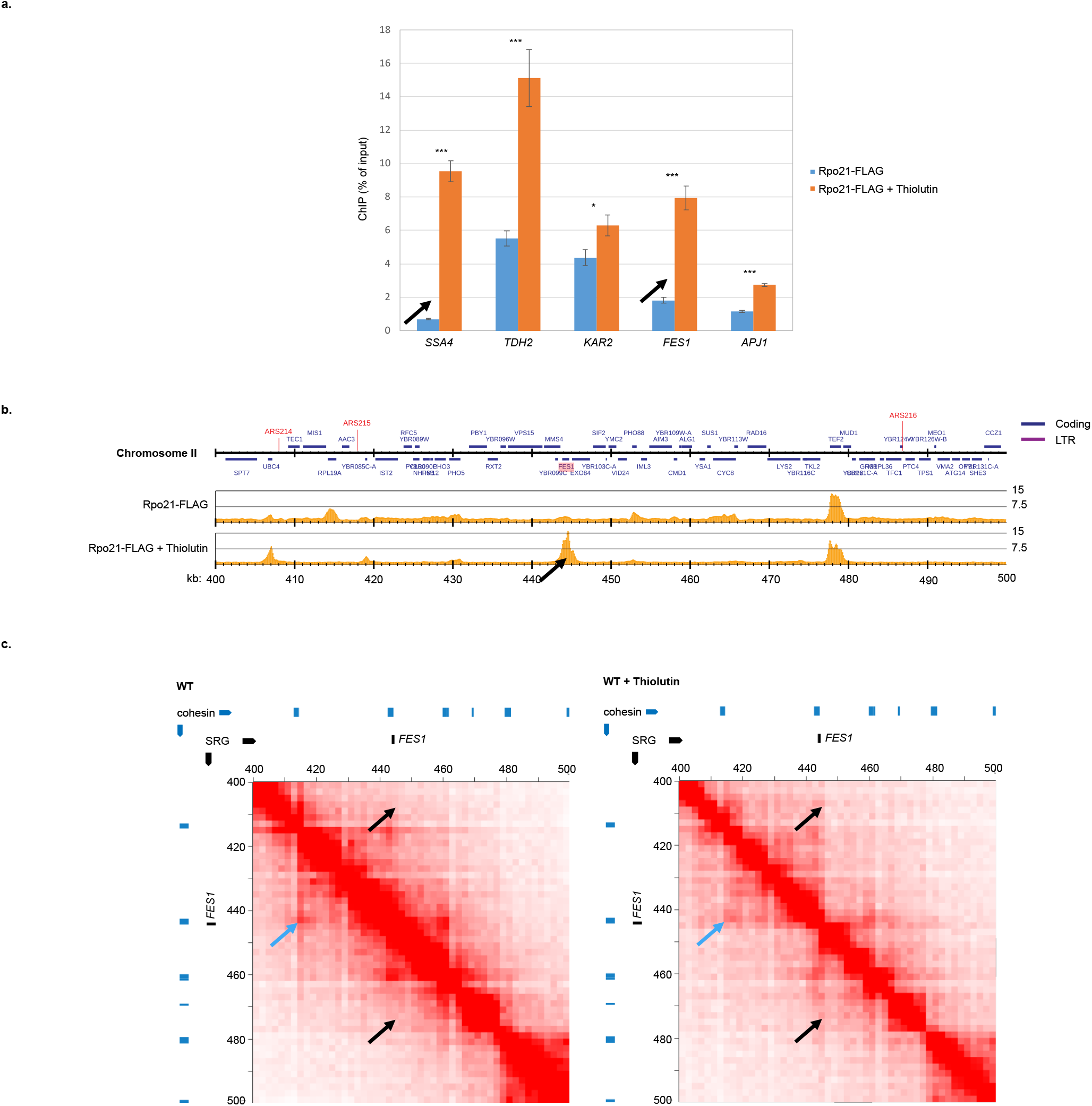
Highly expressed genes create novel chromosome loop barriers. **a.** Chromosome association of Rpo21-FLAG in indicated ORFs as revealed by ChIP-qPCR. Black arrows indicate the barrier-forming genes displayed in c) and Figure 3b. N = 3, ns: p>0.05, *: p≤0.05, ***: p≤0.001. **b.** Map based on normalized ChIP-seq analysis showing Rpo21-FLAG enrichment on part of chromosome II (400-500 kb from left telomere) in the absence or presence of thiolutin, annotations as in Figure 2. Black arrow indicates thiolutin-induced Rpo21-FLAG accumulation at *FES1.* The experiments shown in a) and b) were performed using the growth protocol outlined in Figure S3c. **c.** Normalized Hi-C contact maps (2 kb binning) showing *cis* interactions along the arm of chromosome II (400-500 kb from left telomere) in G2/M-arrested untreated and thiolutin-treated wild type (WT) cells. Black arrows indicate barrier formation and *cis* interactions at the *FES1* gene, induced by thiolutin. Blue arrows highlight interactions that are reduced by thiolutin treatment. Blue and black lines on top and to the left of the panels: cohesin binding sites and stress response gene (SRG), respectively. The experiment in c) was performed using the growth protocol outlined in Figure S3a.

**Figure S7, related to Figure 4.**
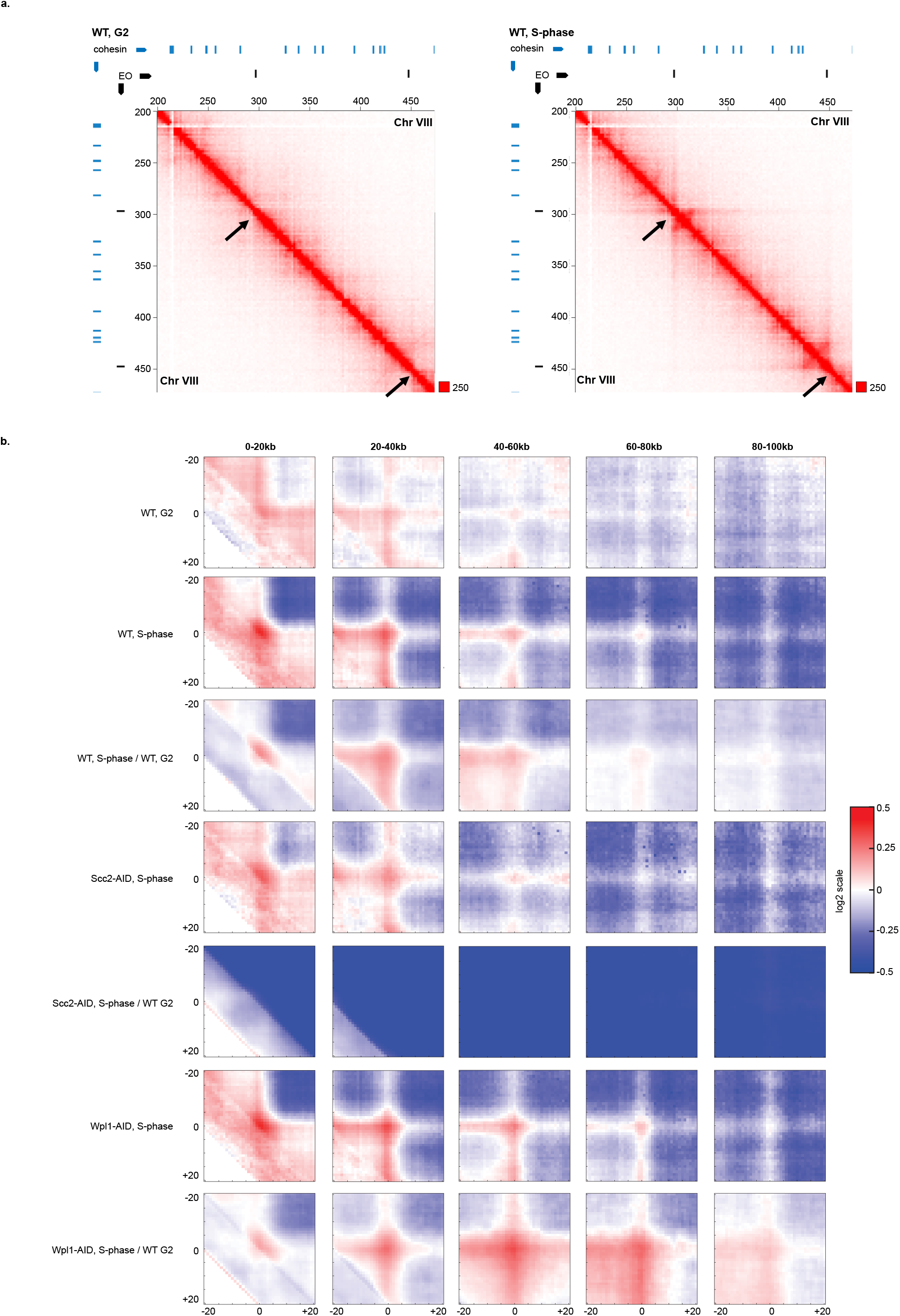
Genome-wide analysis of chromosome loop barriers at replication forks. **a.** Normalized Hi-C contact maps (2 kb binning) showing *cis* interactions along the arm of chromosome VIII (200-450 kb from left telomere) in G2/M- and S-phase-arrested wild type (WT) cells. Black arrows highlight loop boundaries formed at early origins. Blue and black lines on top and to the left of the panels: cohesin binding sites and early firing origin (EO), respectively. **b**. Aggregate peak analysis centered on interactions between pairs of early origins and cohesin sites, separated by increasing chromosomal distances as indicated on top of the panels. The analysis is based on Hi-C analysis of G2/M-arrested WT cells, and S-phase-arrested WT, Scc2- and Wpl1-depleted cells (Scc2-AID, Wpl1-AID). Obtained interaction patterns were either normalized to random sites (e.g. WT, G2) or to origin-cohesin interactions obtained in G2/M-arrested WT cells (e.g. WT, S-phase / WT, G2). Log2 color scale on the right-hand side. The experiments were performed using the growth protocol outlined in Figure S3c (G2/M) cells, and S3d (S-phase).

**Figure S8, related to Figures 1a and 3b.**
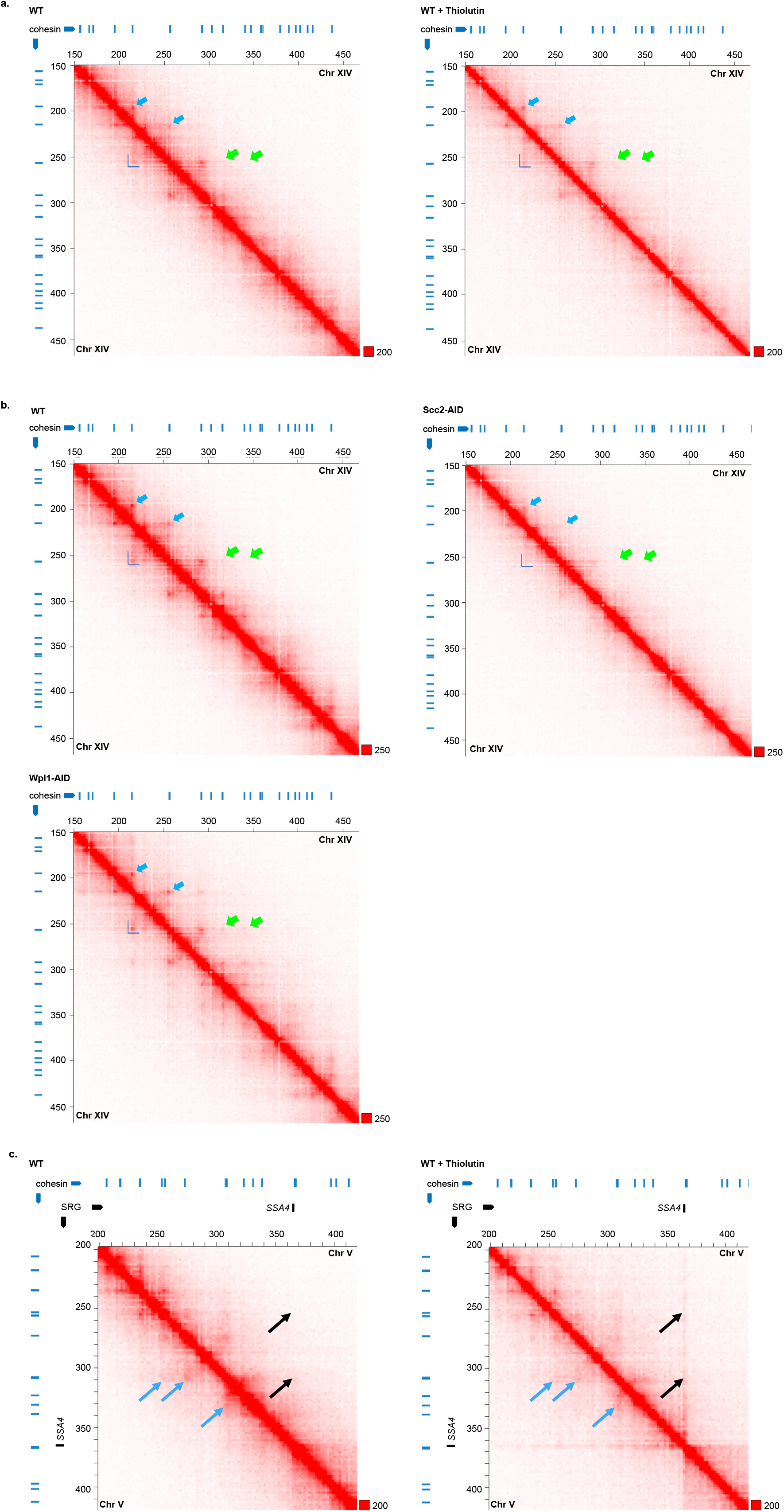
Chromosome *cis* interactions in untreated and thiolutin-treated wild type cells, and after depletion of Scc2 or Wpl1. **a**. Normalized Hi-C contact maps (2 kb binning) showing *cis* interactions along the arm of chromosome XIV (150-450 kb from left telomere) in G2/M-arrested in untreated or thiolutin-treated wild type (WT) cells as indicated. Dark blue L-shape: TAD-like structure, light blue arrows: chromosome loop anchors, light green arrow: chromosome loop anchors detected after Wpl 1 depletion. In contrast to conditions used for Figure 1a, cells were treated with DMSO and thiolutin for 30 minutes after G2/M-arrest, without prior addition of auxin and doxycycline. The experiment was performed using the growth protocol outlined in Figure S3c. **b**. Normalized Hi-C contact maps (2 kb binning) showing *cis* interactions along the arm of chromosome XIV (150-450 kb from left telomere) in G2/M-arrested in wild type (WT) cells, or after Scc2- and Wpl1-depletion (Scc2-AID, Wpl1-AID) as indicated. In contrast to conditions used for Figure 1a, cells were treated with auxin and doxycycline for 1 h after G2/M-arrest without addition of DMSO. In a) and b): Blue lines on top and to the left of the panels: cohesin binding sites, dark blue L-shape: TAD-like structure, light blue arrows: chromosome loop anchors, light green arrow: chromosome loop anchors detected after Wpl1 depletion. **c**. Normalized Hi-C contact maps (2 kb binning) showing *cis* interactions along the arm of chromosome V (200-420 kb from left telomere) in G2/M-arrested untreated and thiolutin-treated wild type (WT) cells. In contrast to conditions used for Figure 3b cells were treated with DMSO and thiolutin for 30 minutes after G2/M-arrest, without prior addition of auxin and doxycycline. Black arrows indicate barrier formation and *cis* interactions at the *SSA4* gene. Blue arrows highlight interactions that are reduced by thiolutin treatment. Blue and black lines on top and to the left of the panels: cohesin binding sites and stress response gene (SRG), respectively. The experiment was performed using the growth protocol outlined in Figure S3c.

**Table S1.** *Saccharomyces cerevisiae* strains used in this study. All strains are of W303 origin *(ade2-1 trp1-1 can1-100 leu2-3,112 his3-11,15 ura3-1) RAD5,* with the modifications listed below.

|  |  |
| --- | --- |
| CB67 | *MATa* |
| CB234 | *MATa SCC1-3xFLAG-KAN* |
| CB1958 | *MATa RPO21-3xFLAG-KAN* |
| CB3060 | *MATa ura3-1::pADH1-OsTIR1-9xMYC-URA3 leu2-3,112::tetR'-SSN6-LEU2* |
| CB3537 | *MATa NAT-tTA-tetO7-3xHA-AID\*-SCC2 ura3-1::pADH1-OsTIR1-9xMYC-URA3 leu2-3,112::tetR'-SSN6-LEU2* |
| CB3593 | *MATa NAT-tTA-tetO7-3xHA-AID\*-WPL1 ura3-1::pADH1-OsTIR1-9xMYC-URA3 leu2-3,112::tetR'-SSN6-LEU2* |

**Table S2.**
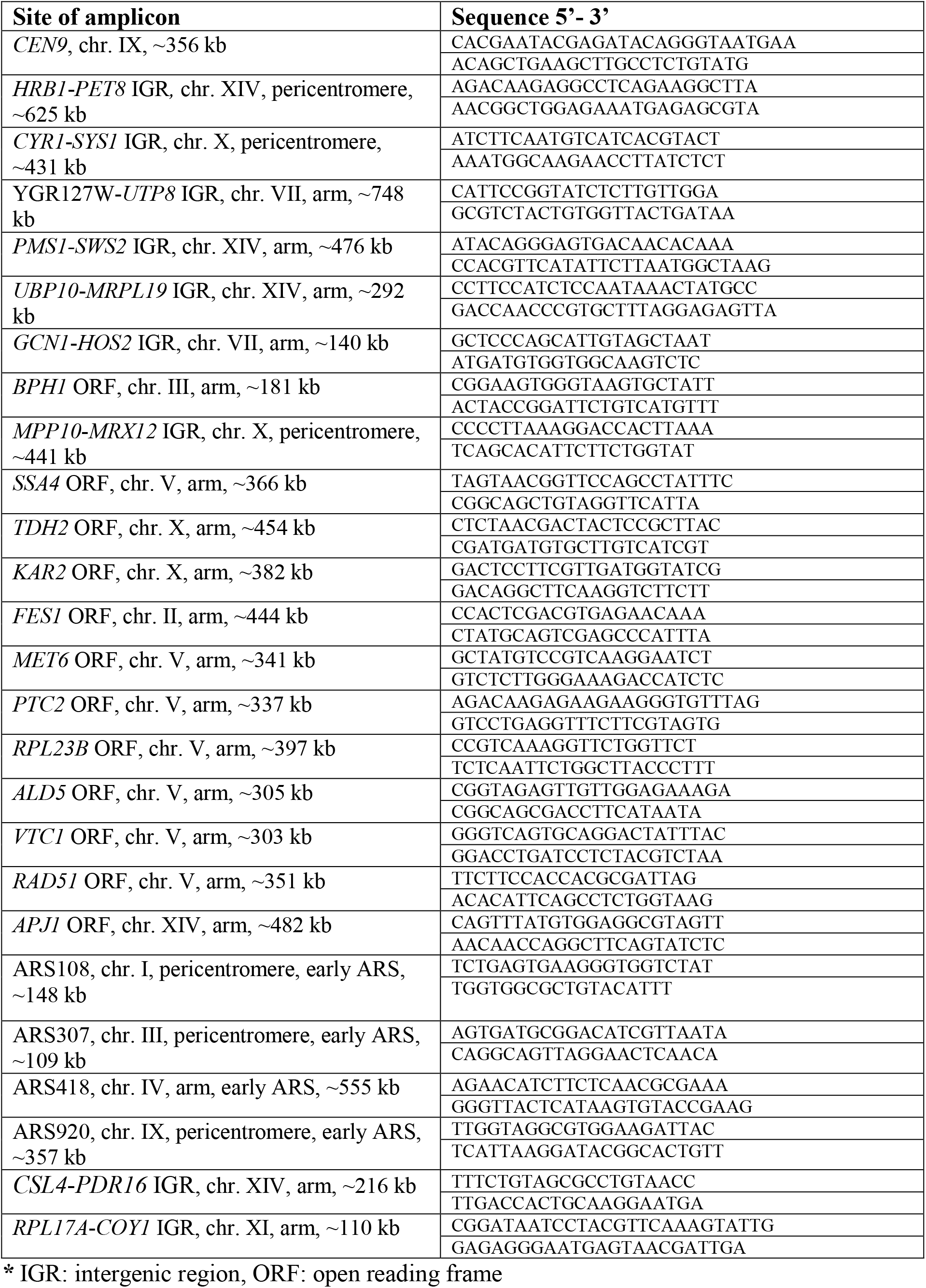
ChlP-qPCR primers

**Table S3.**
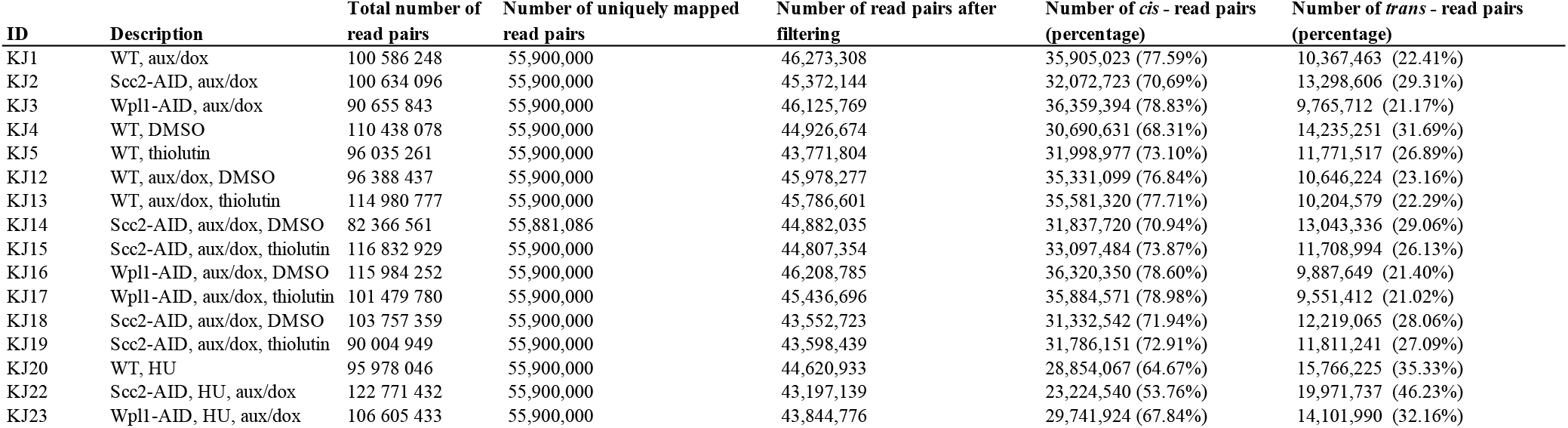
Hi-C statistics

**Table S4.**
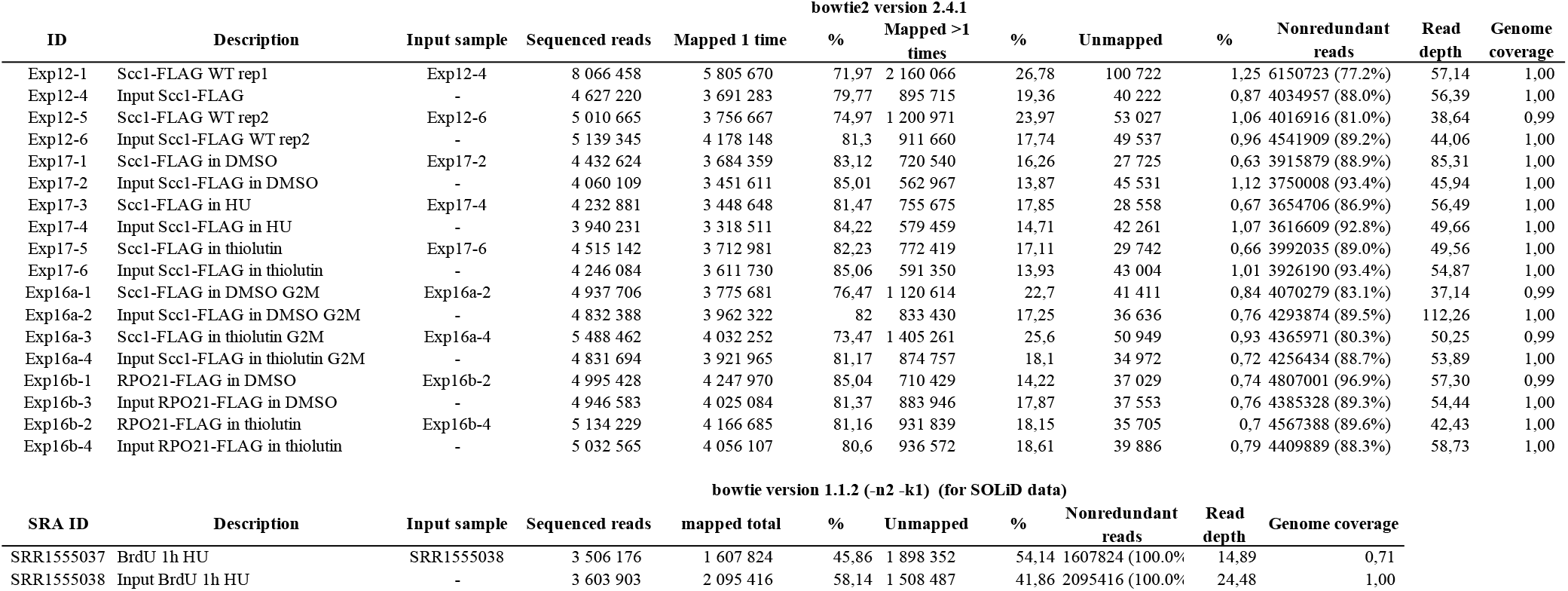
ChIP-seq statistics

| bowtie2 version 2.4.1 |  |  |  |  |  |  |  |  |  |  |  |  |
| --- | --- | --- | --- | --- | --- | --- | --- | --- | --- | --- | --- | --- |
| ID | Description | Input sample | Sequenced reads | Mapped 1 time | % | Mapped >1 times | % | Unmapped | % | Nonredundant reads | Read depth | Genome coverage |
| Exp12-1 | Scc1-FLAG WT rep1 | Exp12-4 | 8 066 458 | 5 805 670 | 71,97 | 2 160 066 | 26,78 | 100 722 | 1,25 | 6150723 (77.2%) | 57,14 | 1,00 |
| Exp12-4 | Input Scc1-FLAG | - | 4 627 220 | 3 691 283 | 79,77 | 895 715 | 19,36 | 40 222 | 0,87 | 4034957 (88.0%) | 56,39 | 1,00 |
| Exp12-5 | Scc1-FLAG WT rep2 | Exp12-6 | 5 010 665 | 3 756 667 | 74,97 | 1 200 971 | 23,97 | 53 027 | 1,06 | 4016916 (81.0%) | 38,64 | 0,99 |
| Exp12-6 | Input Scc1-FLAG WT rep2 | - | 5 139 345 | 4 178 148 | 81,3 | 911 660 | 17,74 | 49 537 | 0,96 | 4541909 (89.2%) | 44,06 | 1,00 |
| Exp17-1 | Scc1-FLAG in DMSO | Exp17-2 | 4 432 624 | 3 684 359 | 83,12 | 720 540 | 16,26 | 27 725 | 0,63 | 3915879 (88.9%) | 85,31 | 1,00 |
| Exp17-2 | Input Scc1-FLAG in DMSO | - | 4 060 109 | 3 451 611 | 85,01 | 562 967 | 13,87 | 45 531 | 1,12 | 3750008 (93.4%) | 45,94 | 1,00 |
| Exp17-3 | Scc1-FLAG in HU | Exp17-4 | 4 232 881 | 3 448 648 | 81,47 | 755 675 | 17,85 | 28 558 | 0,67 | 3654706 (86.9%) | 56,49 | 1,00 |
| Exp17-4 | Input Scc1-FLAG in HU | - | 3 940 231 | 3 318 511 | 84,22 | 579 459 | 14,71 | 42 261 | 1,07 | 3616609 (92.8%) | 49,66 | 1,00 |
| Exp17-5 | Scc1-FLAG in thiolutin | Exp17-6 | 4 515 142 | 3 712 981 | 82,23 | 772 419 | 17,11 | 29 742 | 0,66 | 3992035 (89.0%) | 49,56 | 1,00 |
| Exp17-6 | Input Scc1-FLAG in thiolutin | - | 4 246 084 | 3 611 730 | 85,06 | 591 350 | 13,93 | 43 004 | 1,01 | 3926190 (93.4%) | 54,87 | 1,00 |
| Exp16a-1 | Scc1-FLAG in DMSO G2M | Exp16a-2 | 4 937 706 | 3 775 681 | 76,47 | 1 120 614 | 22,7 | 41 411 | 0,84 | 4070279 (83.1%) | 37,14 | 0,99 |
| Exp16a-2 | Input Scc1-FLAG in DMSO G2M | - | 4 832 388 | 3 962 322 | 82 | 833 430 | 17,25 | 36 636 | 0,76 | 4293874 (89.5%) | 112,26 | 1,00 |
| Exp16a-3 | Scc1-FLAG in thiolutin G2M | Exp16a-4 | 5 488 462 | 4 032 252 | 73,47 | 1 405 261 | 25,6 | 50 949 | 0,93 | 4365971 (80.3%) | 50,25 | 0,99 |
| Exp16a-4 | Input Scc1-FLAG in thiolutin G2M | - | 4 831 694 | 3 921 965 | 81,17 | 874 757 | 18,1 | 34 972 | 0,72 | 4256434 (88.7%) | 53,89 | 1,00 |
| Exp16b-1 | RPO21-FLAG in DMSO | Exp16b-2 | 4 995 428 | 4 247 970 | 85,04 | 710 429 | 14,22 | 37 029 | 0,74 | 4807001 (96.9%) | 57,30 | 0,99 |
| Exp16b-3 | Input RPO21-FLAG in DMSO | - | 4 946 583 | 4 025 084 | 81,37 | 883 946 | 17,87 | 37 553 | 0,76 | 4385328 (89.3%) | 54,44 | 1,00 |
| Exp16b-2 | RPO21-FLAG in thiolutin | Exp16b-4 | 5 134 229 | 4 166 685 | 81,16 | 931 839 | 18,15 | 35 705 | 0,7 | 4567388 (89.6%) | 42,43 | 1,00 |
| Exp16b-4 | Input RPO21-FLAG in thiolutin | - | 5 032 565 | 4 056 107 | 80,6 | 936 572 | 18,61 | 39 886 | 0,79 | 4409889 (88.3%) | 58,73 | 1,00 |
| bowtie version 1.1.2 (-n2 -k1) (for SOLiD data) |  |  |  |  |  |  |  |  |  |  |  |  |
| SRA ID | Description | Input sample | Sequenced reads | mapped total | % | Unmapped | % | Nonredundant reads | Read depth | Genome coverage |  |  |
| SRR1555037 | BrdU 1h HU | SRR1555038 | 3 506 176 | 1 607 824 | 45,86 | 1 898 352 | 54,14 | 1607824 (100.0%) | 14,89 | 0,71 |  |  |
| SRR1555038 | Input BrdU 1h HU | - | 3 603 903 | 2 095 416 | 58,14 | 1 508 487 | 41,86 | 2095416 (100.0%) | 24,48 | 1,00 |  |  |

**Table S5.**
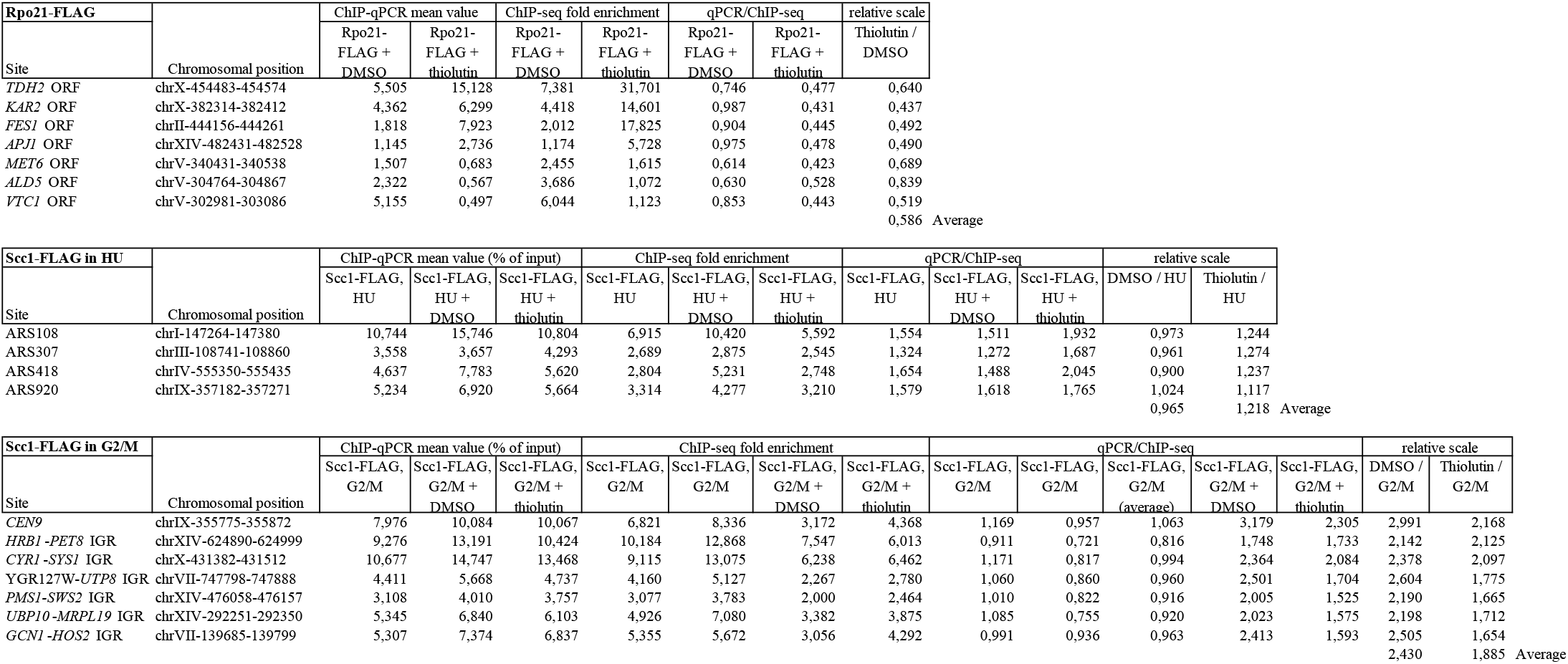
ChlP-seq ChlP-qPCR normalization

